# Deep transfer learning-based hologram classification for molecular diagnostics

**DOI:** 10.1101/192559

**Authors:** Sung-Jin Kim, Chuangqi Wang, Bing Zhao, Hyungsoon Im, Jouha Min, Hee June Choi, Joseph Tadros, Nu Ri Choi, Cesar M. Castro, Ralph Weissleder, Hakho Lee, Kwonmoo Lee

## Abstract

Lens-free digital in-line holography (LDIH) is a promising microscopic tool that overcomes several drawbacks (e.g., limited field of view) of traditional lens-based microcopy. However, extensive computation is required to reconstruct object images from the complex diffraction patterns produced by LDIH, which limits LDIH utility for point-of-care applications, particularly in resource limited settings. Here, we describe a deep transfer learning (DTL) based approach to process LDIH images in the context of cellular analyses. Specifically, we captured holograms of cells labeled with molecular-specific microbeads and trained neural networks to classify these holograms without reconstruction. Using raw holograms as input, the trained networks were able to classify individual cells according to the number of cell-bound microbeads. The DTL-based approach including a VGG19 pretrained network showed robust performance with experimental data. Combined with the developed DTL approach, LDIH could be realized as a low-cost, portable tool for point-of-care diagnostics.

## INTRODUCTION

Lens-free digital in-line holography (LDIH) is a powerful imaging platform that overcomes many of the limitations of traditional microscopy^1–6^. LDIH records diffraction patterns produced by samples, which can later be used to computationally reconstruct original object images. This strategy enables LDIH to image a large area (~mm^2^) while achieving a high spatial resolution (~µm). Furthermore, the simplistic optical design allows for compact setups, consisting of a semiconductor imager chip and a coherent light source. LDIH has been previously tested for potential point-of-care (POC) diagnoses^7^. Recently, we have advanced LDIH for the purpose of molecular diagnostics (D3, digital diffraction diagnostics)^3^ wherein cancer cells were labeled with antibody-coated-microbeads, and bead-bound cells were counted for molecular profiling.

A major hurdle to translating LDIH into POC tests is the need for extensive computational power. In principle, diffraction patterns can be back-propagated to reconstruct human-friendly object images. The bottleneck lies in the recovery of phase information, lost during the imaging process. It has been shown that this information can be numerically recovered through iterative optimization^1,8–13^, but the process is costly in computation time and requires high-end resources. To overcome this issue, it was demonstrated that a deep neural network could be trained to recover phase information and reconstruct object images, substantially reducing the total computational time^14^. However, this method still required an input of back-propagation images obtained from the holograms. In this paper, we explored an alternative approach in which diagnostic information could be extracted from the raw hologram images without the need for hologram reconstruction. In the microbead-based assay, we reasoned that cell-bead objects could generate distinct hologram patterns, albeit imperceptible to human eyes, recognizable by machine vision classifiers. Developing such a capacity would eliminate the need for image reconstruction, further advancing LDIH utility for POC operations.

We here report on new machine-learning (ML) based approaches for LDIH image analysis. ML has been making significant progress in extracting information from complex biomedical images and started to outperform human experts for many data sets^15–18^. In this paper, we took deep transfer learning (DTL)^19–25^ approach to classify raw holograms and compared them with other ML schemes including convolutional neural networks (CNN)^26–28^. DTL extracts feature information from input data using the convolution part of pre-trained networks and subsequently feeds the information to classifiers. It has been known that pretrained networks can be exploited as a general-purpose feature extractor^20^. In this DTL approach, we used a VGG19^29^ model that was pretrained with a large number of ordinary images (i.e., not holograms) available in the ImageNet^30^, and fine-tuned the classifier to obtain high-performance classification. We applied these approaches to classifying holograms generated from cells and microbeads without a reconstruction process. Specifically, algorithms were developed to i) automatically detect the holograms of cells labeled with microbeads, ii) classify detected cells according to the number of the cell-bound beads, and iii) construct the histogram of the cell-bound beads from the entire hologram. We found that a DTL approach offered more reliable, robust, and efficient performance in hologram classification than the conventional CNN.

## RESULTS

### System and assay setup

**Figure 1A** shows the schematic of LDIH system^3^. As a light source, we used a light-emitting diode (LED; λ = 420 nm). The light passes through a circular aperture (diameter, 100 μm), generating a coherent spherical wave on the sample plane. The incidence light and the scattered light from the sample interfere with each other to generate holograms which are then recorded by a CMOS imager^10,31^. The system has a unit (×1) optical magnification, resulting in a field-of-view equal to the imager size.

To enable molecular-specific cell detection, we used antibody-coated microbeads (diameter, 6 μm) for cell labeling. The number of attached beads is proportional to the expression level of a target marker, allowing for quantitative molecular profiling^3^. Diffraction patterns from unlabeled and bead-bound cells have subtle differences that are hard to detect with human eyes (**Fig. 1B**). Only after image reconstruction can beads and cells be differentiated and counted; cells have high amplitude and phase values, whereas microbeads have negligible phase values.

**Figure 1.**
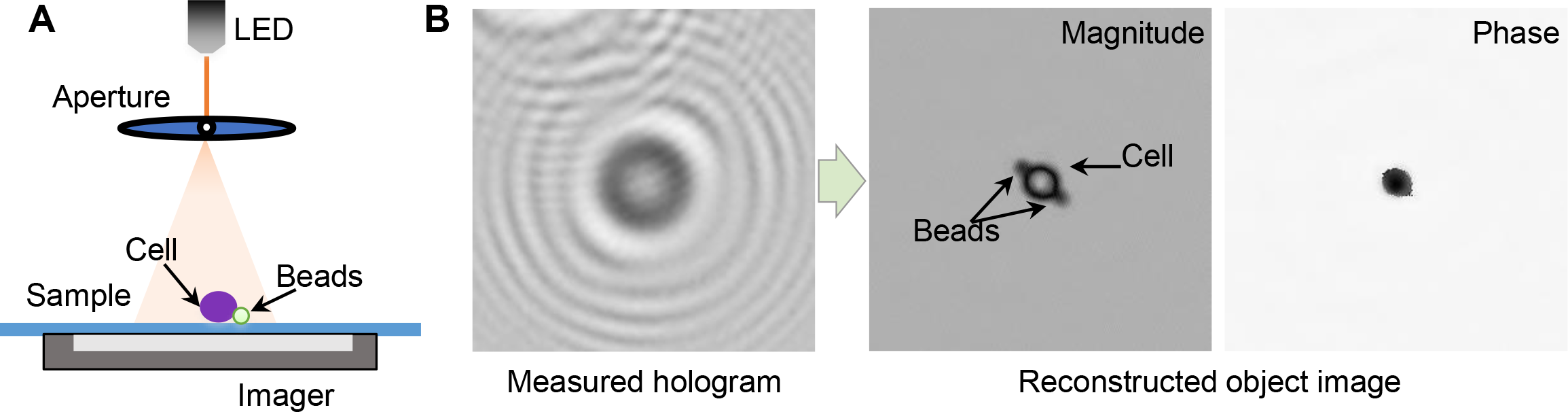
In-line holographic imaging. **(A)** A holography system includes LED, a sample glass and a sensor where light is passed to a sample through a pinhole disk. **(B)** A hologram image and its associated reconstructed images consisting of magnitude and phase images.

### Reconstruction-free ML approaches

Conventional LDIH reconstruction (**Fig. 2A**) requires multiple repetitions of back-propagation, constraint application, and transformation^8^. This iterative algorithm is computationally intensive, either incurring long processing time or requiring high-end resources (e.g., a high-performance graphical processing unit server) for faster results^3^. Furthermore, human curation is occasionally needed to correct for stray reconstruction (e.g., debris, twin images). In contrast, our ML-based approach is a reconstruction-free classification method (**Fig. 2B**). As an off-line task, we first build a training dataset by automatically detecting cell candidates and cropping them from the entire holograms. Then, we labeled each cropped hologram with the number of the cell-bound beads using reconstructed image as ground truth. Next, we trained a network using annotated holograms of bead-bound cells. After the training was complete, the network was used for on-line classification tasks; cell candidate holograms were detected and their holograms, without any image preprocessing, were entered as input for classification based on the number of cell bound beads. Finally, the histograms of the cell-bound beads from the entire holograms were created for molecular diagnosis.

**Figure 2.**
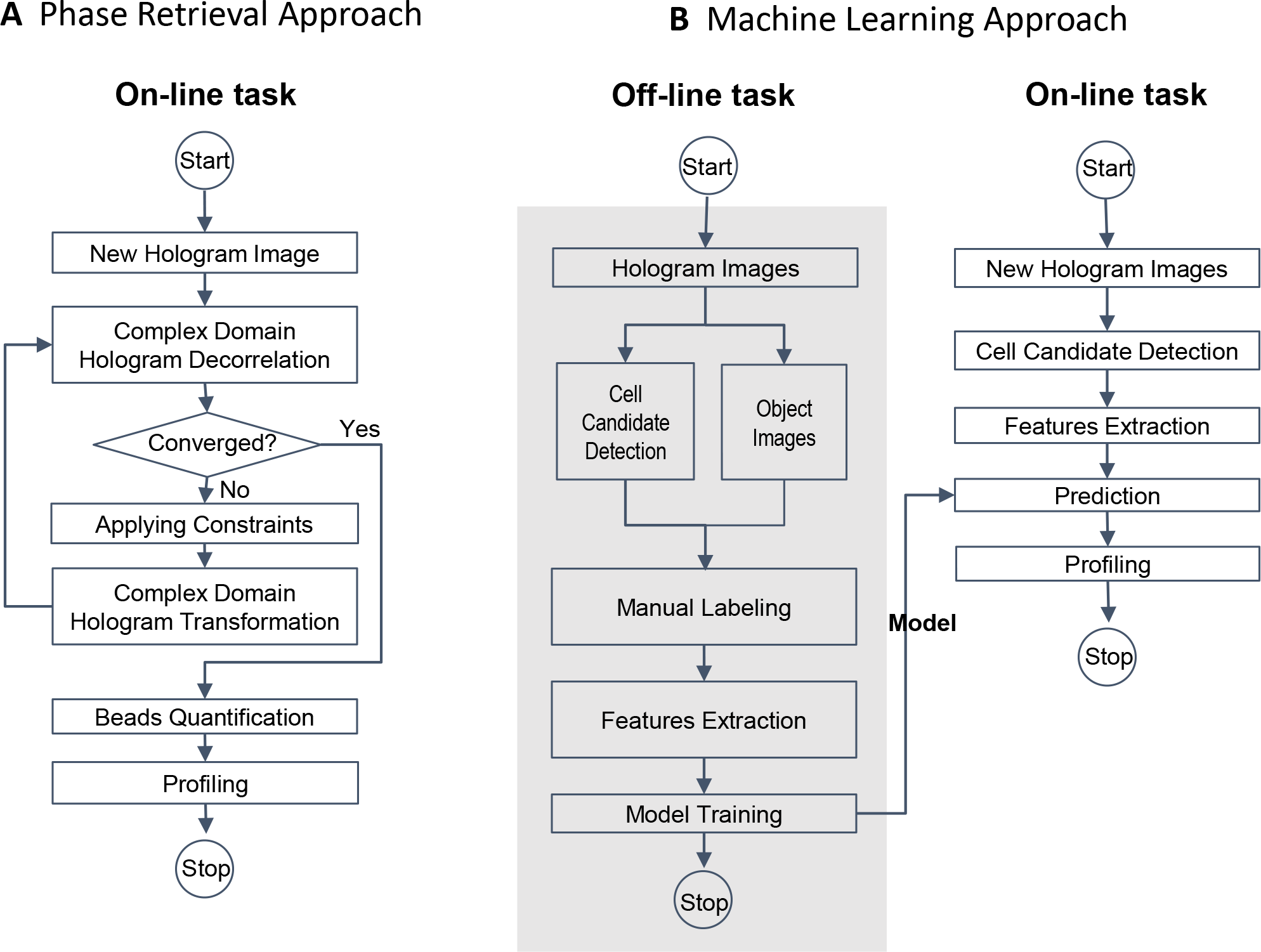
Flow charts of holographic diagnostic approaches. **(A)** A conventional approach includes iterative reconstruction processes by phase retrieval. **(B)** A machine learning based workflow for hologram classification.

Both off-line and on-line tasks in the ML approach (Fig. 2B) required the automatic detection of holographic patterns of cells. To achieve this task, we implemented a computational method which identifies the center of individual diffraction patterns^32^. First, the images of the gradient magnitude of holograms were generated and thresholded based on their intensity. The converging directions of gradients were used to estimate the positions of cell candidates in holograms (**Fig. 3A**; see Methods). Using this method, we detected 2729 potential cell candidates from 31 holograms. The samples for these holograms were prepared by labeling SkBr3 breast carcinoma cells with polystyrene beads conjugated with control, EpCAM, and HER2 antibodies. Then, we reconstructed object images and cropped the holograms and the object images (270 × 270 pixels). We labeled the cropped holograms **(Fig. 3C**) and their reconstructed object images (**Fig. 3D**) with the number of the bead attached to a cell (*N*_*B*_ : 0, 1, 2, 3, ≥ 4). There were also images of floating beads, multiple cells, and artifacts, which were collectively labeled as ‘background’ (BG). The distribution of the class in the final training set is shown in **Fig. 3B**.

**Figure 3.**
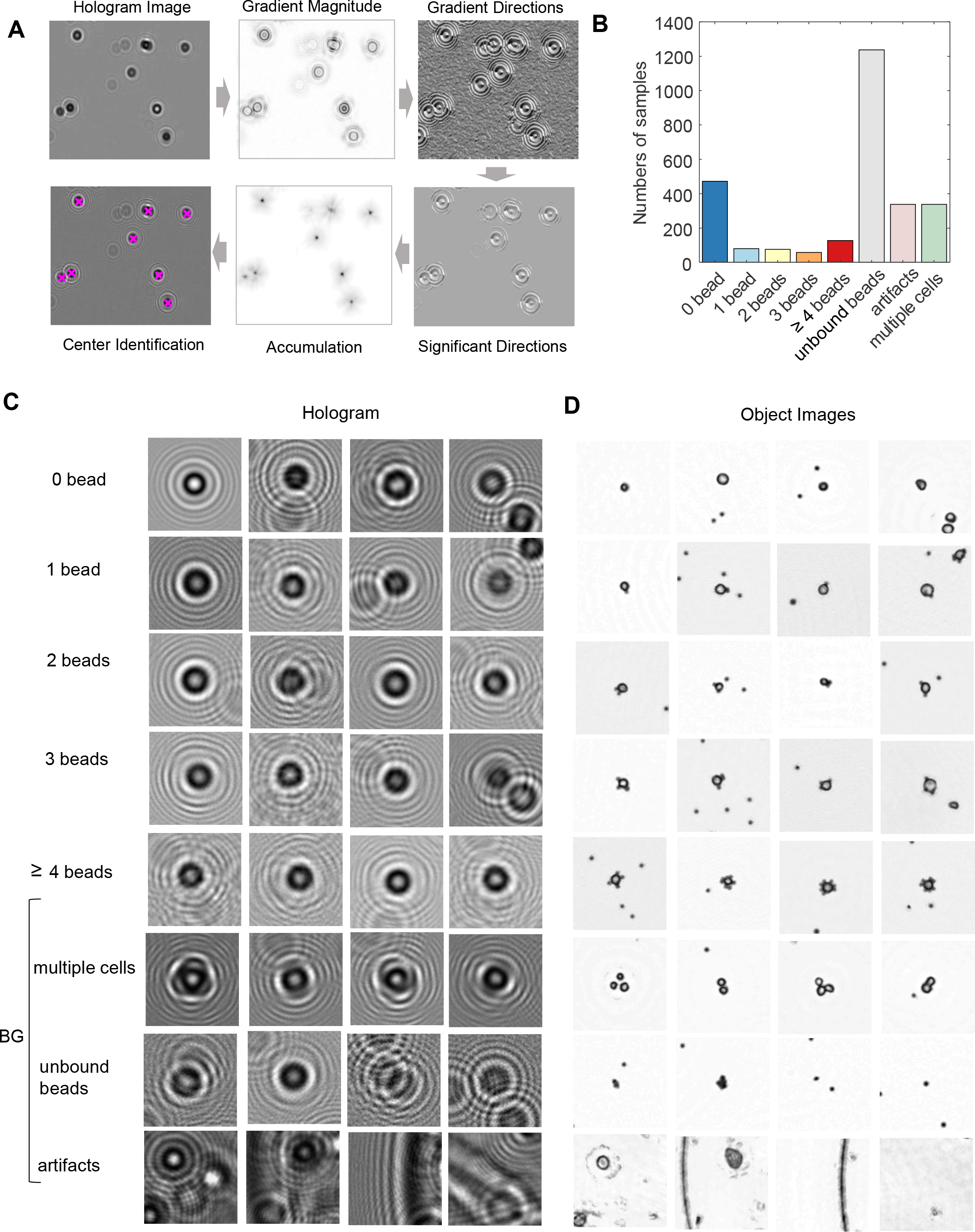
Training set preparation for hologram classification. **(A)**Cell candidates detection workflow. **(B)**Class distribution in the training dataset. **(C-D)**Sample examples of holograms (C) and corresponding object images (D).

### Visualization of hologram features

We first tested the feasibility of the reconstruction-free classification by visualizing the features extracted from the holograms. Using VGG19 pretrained model, we extracted features from the training set of holograms (**Fig. 4A**). Since VGG19 was trained using color images (RGB channels) and our data were in a gray scale, we copied the same image in each channel in the VGG-19 pretrained model. Then, using PCA (Principal Component Analysis), we reduced the feature dimension from 32,768 to 500 and visualized their distribution using t-SNE plots^33^. In both holograms (**Fig. 4B**) and object images (**Fig. 4C**), each class of bead-bound cells was visually more segregated than the cases where only the same PCA was applied without using VGG19 (**Fig. 4D-E**), suggesting that VGG19-based features of the holograms could discriminate the numbers of cell-bound beads.

**Figure 4.**
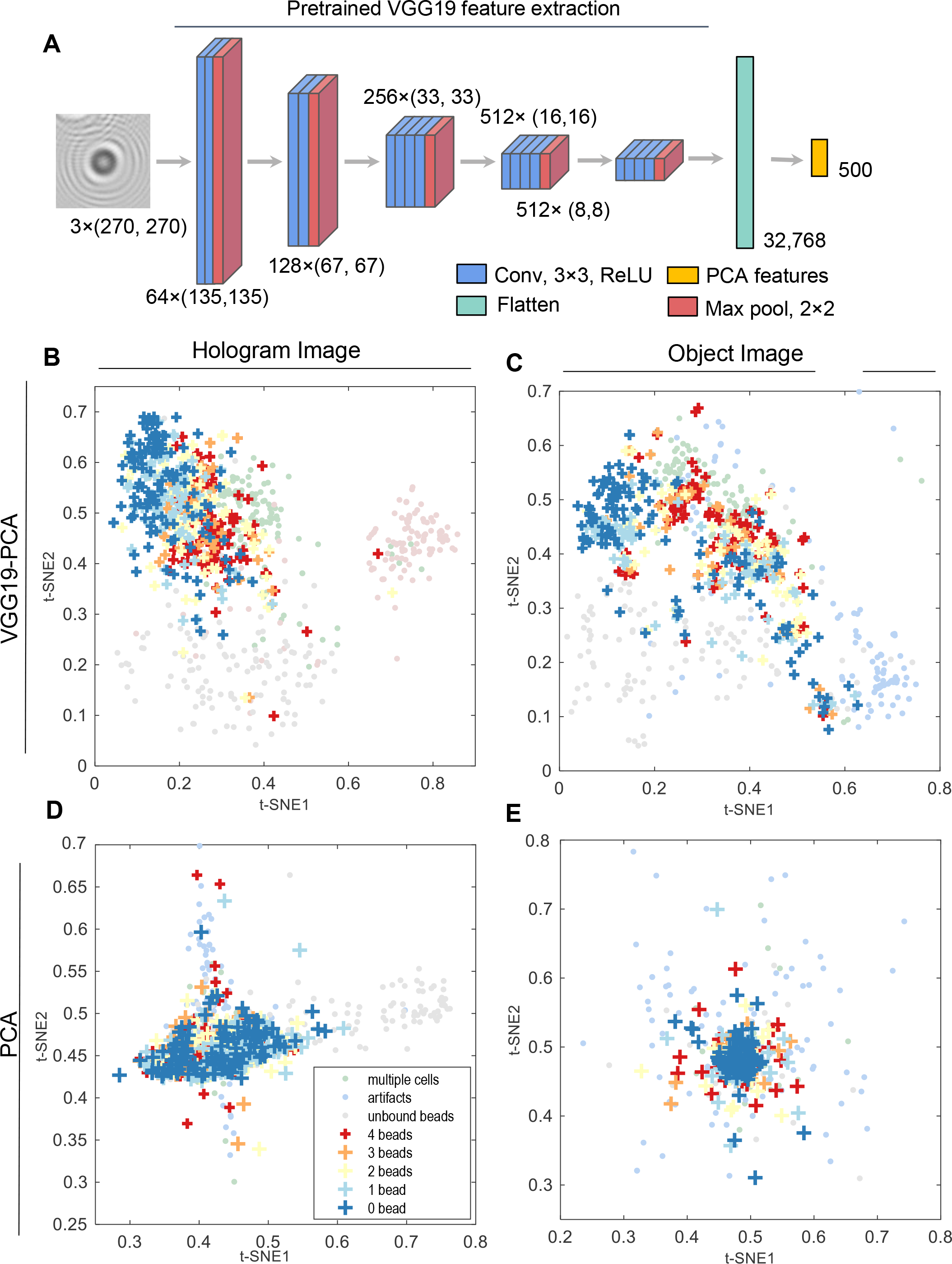
Feature extraction from holograms. **(A)** Features extraction by the pretrained VGG19 model and PCA. **(B-D)** t-SNE plots of the extracted features. VGG19-PCA feature extraction from the holograms (B) and object images (C). PCA feature extraction from the holograms (D) and object images (E)

### Classification results by deep transfer learning

Using the features from VGG19-PCA or PCA, we trained the multilayer perceptron (MLP, **Fig. 5A**) separately for holograms or object images. Since the training data were unbalanced (**Fig. 3B**), we took the following approaches: To balance the training set of cell-bound beads, we applied the data augmentation (rotation and zoom-in) to increase the data size in the case of *N*_*B*_ ≥ 1 (see Methods). Then, to address the unbalance between bead-bound cells (*N*_*B*_ : 0 to ≥ 4) and background (BG) data we used the weighted cost function using the proportion of bead-bound cells to BG data. From the whole dataset consisting of 2729 cropped images from 31 holograms, we randomly split the data into training, validation, and testing dataset with a 64:16:20 ratio. The model was selected based on the validation loss, and the model performance was evaluated based on the testing data. For statistical analysis, we repeated the training 20 times with different data splitting (see Methods for detail).

**Figure 5.**
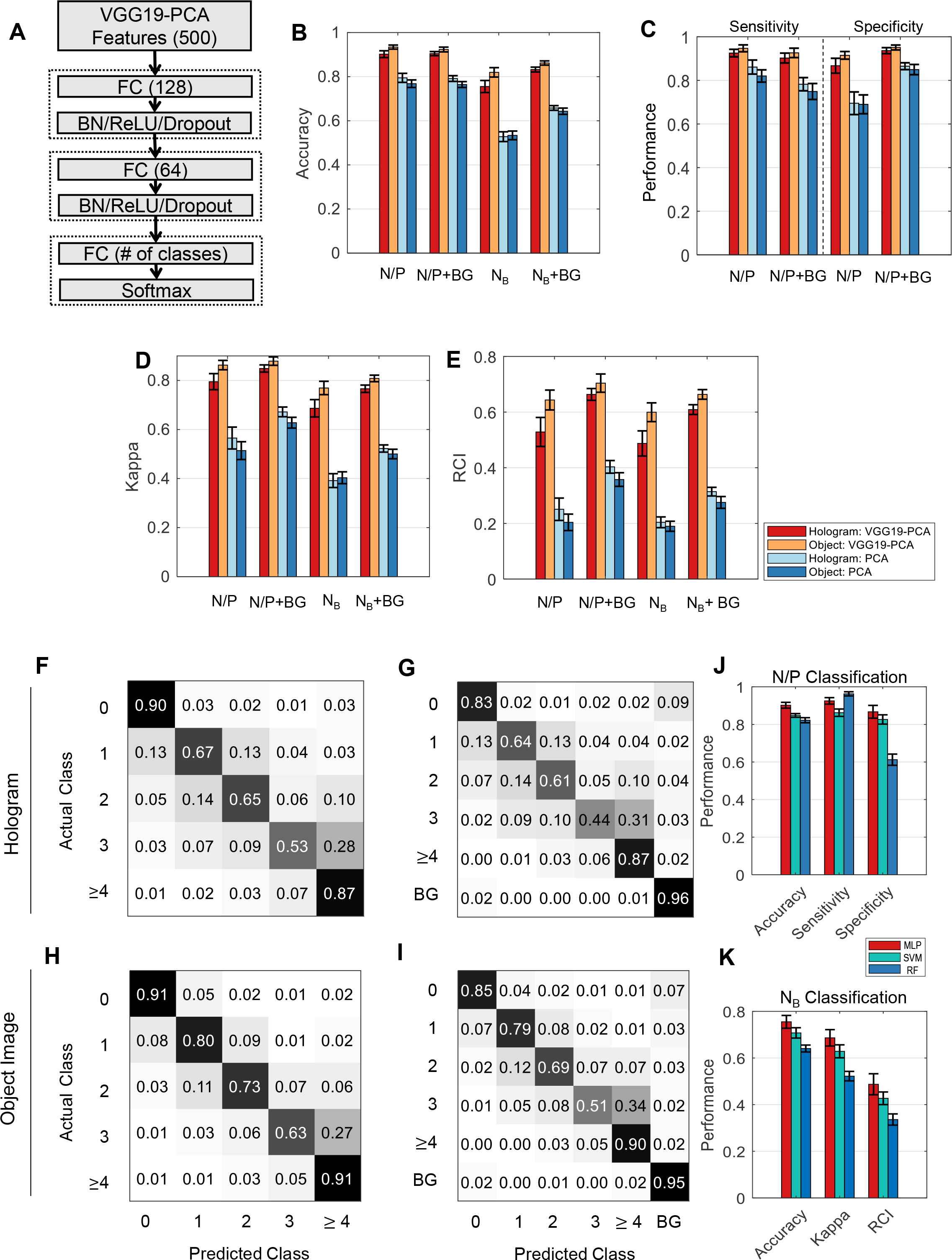
Classification performance of the deep transfer learning for holograms. **(A)** MLP (Multi-Layer Perceptron) neural network classifier used in this study (FC: fully connected layer, BN: batch normalization layer). **(B-E)** Performance Comparison between VGG19-PCA-MLP and PCA-MLP. N/P: negative (*N*_*B*_≤ 1) and positive (*N*_*B*_≥ 2) bead attachment. N/P+BG: *N*_*B*_≤ 1, NB≥ 2, BG (background class), *N*_*B*_: the numbers of beads (0, 1, 2, 3, ≥ 4 beads), *N*_*B*_+BG: the numbers of beads (0, 1, 2, 3, ≥ 4), BG. The performance measures are accuracy (B), sensitivity/specificity (C), Cohen’s Kappa (D), and RCI (E). **(F-K)** Average confusion matrices using VGG19-PCA-MLP using *N*_*B*_ (five classes) classification (F) and NB+BG (6 classes) classification (G) for holograms, and *N*_*B*_ classification (H), *N*_*B*_+BG classification (I) for object images. **(J-K)** The performance comparisons of VGG19-PCA-MLP (MLP), Support Vector Machine (SVM), Random Forest (RF) in N/P (two classes) classification (J) and *N*_*B*_ (five classes) classification (K) using holograms.

Since cells with more than two bead attachments are considered positive for a given target biomarker^3^, we first performed the binary classification (N/P) based on the bead number (*N*_*B*_): negative (*N*_*B*_≤ 1) vs. positive (*N*_*B*_≥ 2). The accuracies of VGG19-PCA-MLP in N/P were 90.2% for holograms and 93.4% for object images, whereas the accuracies of PCA-MLP were only 79.5% for holograms and 76.8% for object images (**Fig. 5B** and **Supplementary Table 1**). Since the background (BG) data are included in real situations, we also trained the classifiers after adding the BG class (N/P+BG). In comparison to the N/P classification, the accuracies were still similar (VGG19-PCA-MLP: 90.4% for holograms and 92.3% for object images, PCA-MLP: 79.1% for holograms and 76.4% for object images) (**Fig. 5B** and **Supplementary Table 1)**. The sensitivity and specificity also showed that VGG19-PCA-MLP outperformed PCA-MLP in all cases (**Fig. 5C, Supplementary Table 2 and 3**).

While this binary classification for the negative and positive cells can be applied to molecular diagnostics, the actual number of the beads and their distribution from a given patient sample can provide more detailed information including cancer stages and patient sub-types. Therefore, we trained the classifiers based on the numbers of the cell-bound beads. When we performed the classification using the number of beads (0, 1, 2, 3, ≥ 4), VGG19-PCA-MLP achieved significantly higher accuracies, 75.5% for holograms and 82.0% for object images than PCA-MLP (52.8% for holograms and 53.4% for object images) (**Fig. 5B** and **Supplementary Table 1**). When BG class was considered together for the real application (*N*_*B*_+BG), VGG19-PCA-MLP achieved 83.2% for holograms, and 86.2% for object images, whereas PCA-MLP achieved 65.8% for holograms and 64.3% for object images (**Fig. 5B** and **Supplementary Table 1)**. The distinctiveness of the BG class from the other classes (**Fig. 4B-C**) increased the overall classification accuracies.

To quantitatively compare the classification performance among all classification cases, we employed the Cohen’s kappa coefficient^34^ and the relative classifier information (RCI) ^35,36^ (**Fig. 5D-E**, **Supplementary Table 4** and **5**). Cohen’s kappa compensates for classifications by random chance, and RCI quantifies how much uncertainty had been reduced by the classification relative to the prior probabilities of each class. Both measures are between 0 and 1 (0: worst, 1: perfect classification). The *N*_*B*_ classification using VGG19-PCA-MLP and the holograms produced the significantly larger values of Cohen’s kappa (0.687) and RCI (0.487) than the PCA-MLP (Cohen’s kappa: 0.392, RCI, 0. 0.204, **Supplementary Table 4** and **5**). The results showed the followings; i) N/P classifiers has better performance than *N*_*B*_ classifiers since multi-category classification is more prone to error than binary classification. ii) The VGG19-PCA-MLP outperforms PCA-MLP in all cases, iii) While the classification using holograms showed good performance, the classification using object images performs marginally better than holograms.

To see the performance of the multi-category classification more closely, we computed the average confusion matrices of the bead classification (*N*_*B*_, *N*_*B*_+BG). The prediction accuracy was high when the number of beads is 0 or ≥4, or the BG class, and the accuracies decreased when the bead number was between 1 and 3 (**Fig. 5F-I**). Since the high occurrence values of the confusion matrices were near the diagonals, the misclassification mainly happened among neighboring numbers for both holograms and object images. This property makes the molecular profiling from the entire holograms less susceptible to mis-classification error.

To compare the performances of different classifiers in our DTL approach, we also trained SVM (Support Vector Machine) and RF (Random Forest) using the same VGG19-PCA features. In both N/P and *N*_*B*_ classifications, MLP outperformed SVM and RF significantly (see p-values in **Supplementary Table 6 and 7**).

### Molecular profiling using the deep transfer learning

When it comes to the molecular diagnosis using LDIH, the clinical decision is often made at the cell population level. Therefore, we assessed how our hologram multi-category classification (*N*_*B*_) matched with the distribution of the cell-bound beads from an entire hologram. We overlaid the actual and predicted distributions of cell-bound beads from 18 different samples, whose number of detected cell candidates are more than 15 excluding BG (**Fig. 6A**). We also plotted that the histograms of the actual and predicted numbers of the cell-bound beads in each sample (**Fig. 6B**). These show that the predicted bead proportions matched well with the actual distribution. Also, the mean difference between the proportions of the actual and the prediction was within 5% (**Fig. 6E**). This suggests that our multi-category classification based on the number of the cell-bound beads can be used to characterize the molecular profiles of the cancer cell population from a patient sample.

**Figure 6.**
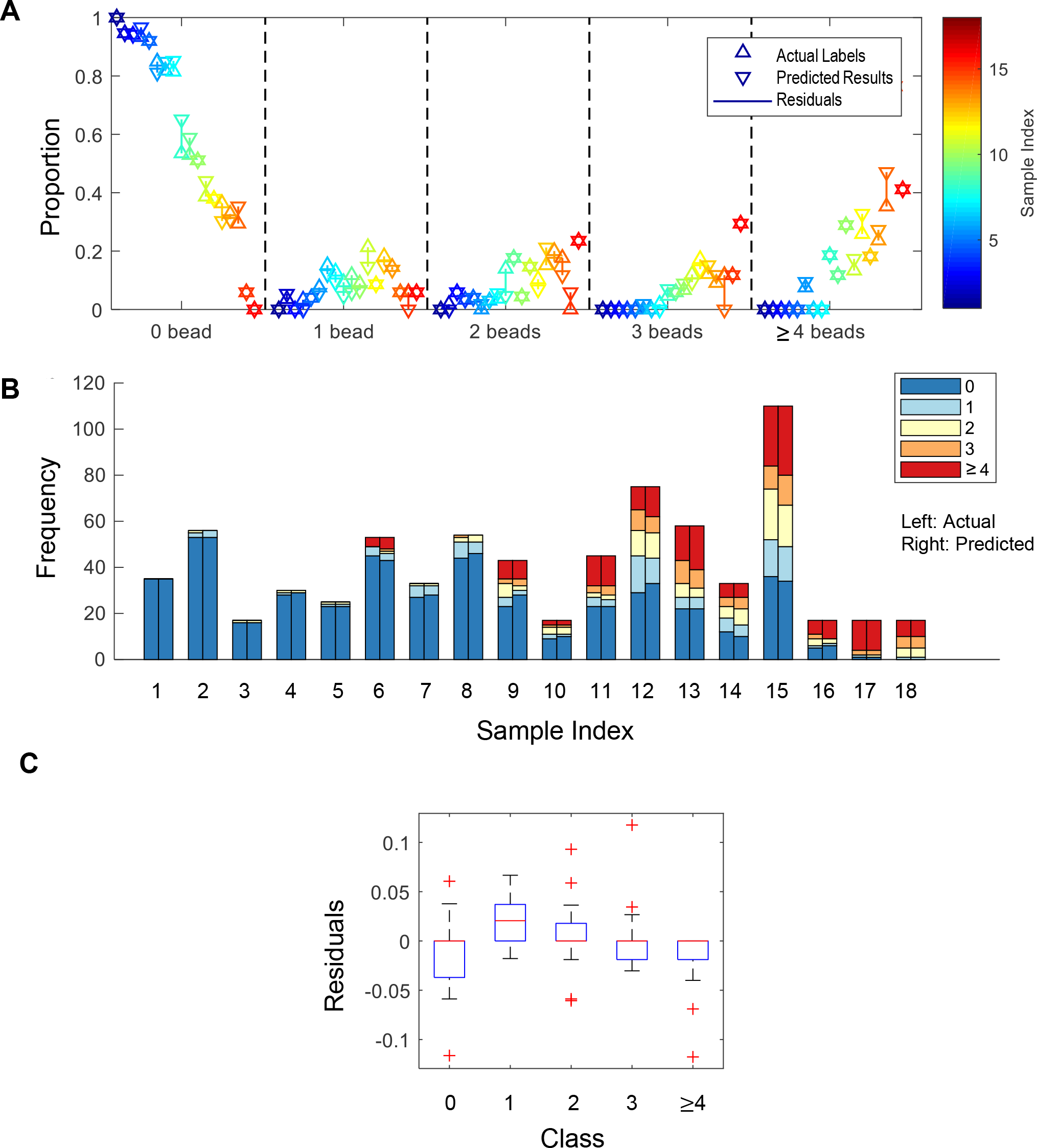
Molecular profiling using deep transfer learning. **(A-B)** Comparison of the proportions (A) and numbers (B) of the cell-bound beads in each entire hologram between the actual labels (left) and predicted results (right). The classifier was built without BG class (*N*_*B*_). The color represents each hologram. **(C)** The residuals between actual and predicted proportions in each class

### Roles of VGG19 pretrained model

To evaluate the role of the pretrained model in our classification, we also trained a conventional convolutional neural network (CNN) *de novo* for *N*_*B*_ classification (**Fig. 7A**). Whereas there were no statistical differences in accuracy, RCI, and kappa between the CNN and the VGG19-PCA-MLP (**Fig. 7B, Supplementary Table 8**), we observed that the validation loss and accuracy of the training curves of this CNN were far more fluctuating than VGG19-PCA-MLP (**Fig. 7C-D**). Therefore, the standard deviation of the accuracy and the loss in the steady state of the CNN training was significantly larger than those of VGG19-PCA-MLP (p-values: 2.30 × 10^−107^ for the loss, and 1.03 × 10^−255^ for the accuracy by two sample F-test) (**Fig. 7E-F**). Since the CNN has much more parameters to learn than VGG19-PCA-MLP, the cost function of the validation set may have much more local minima than that of the training set, which makes the validation loss and accuracy fluctuating during the training process. This suggests that DTL is more robust to the data variability and can produce a more generalizable classifier.

**Figure 7.**
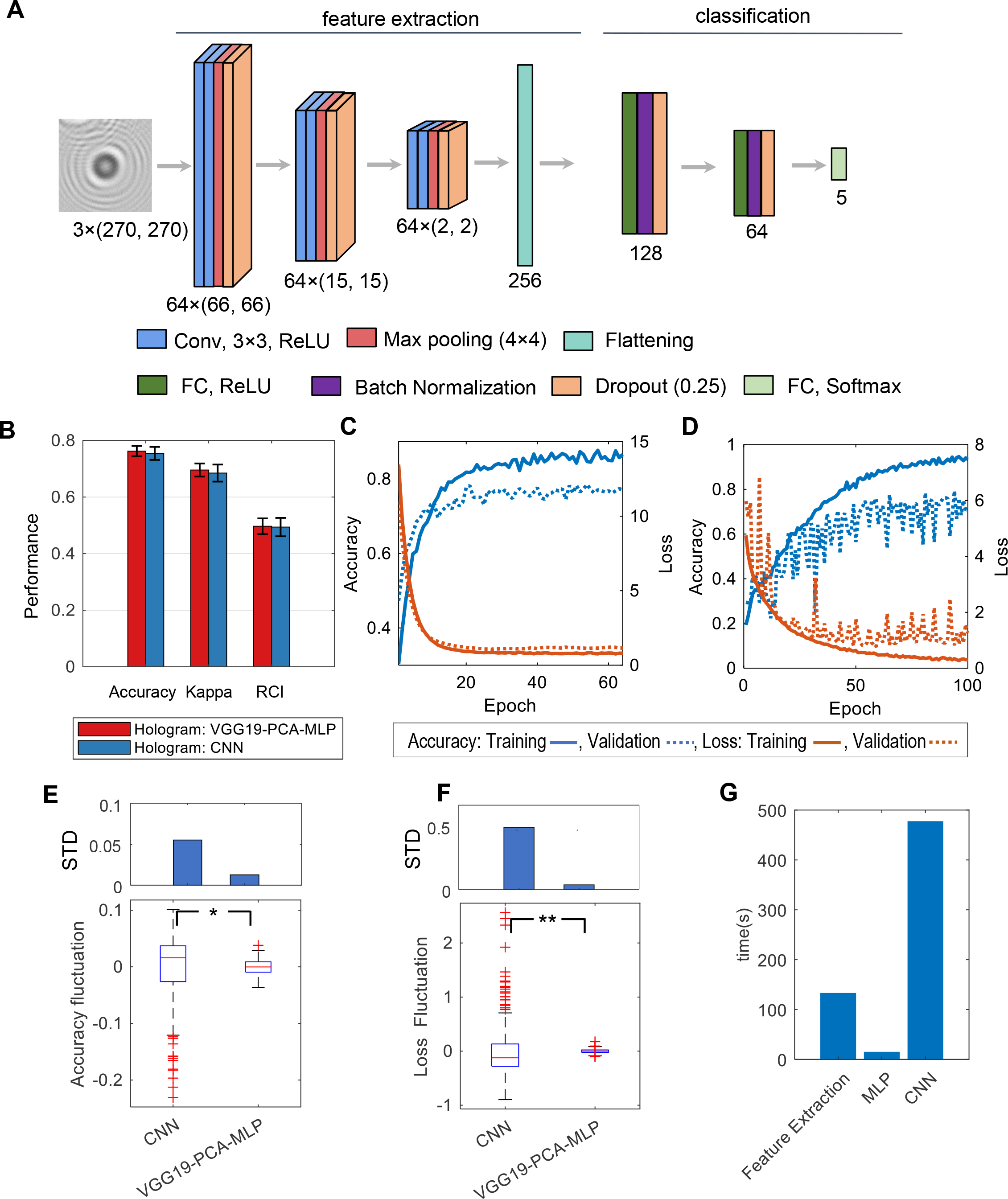
Classification performance of convolution neural networks (CNN). **(A)** The structure of the CNN used in this study. **(B)** Performance comparison between VGG19-PCA-MLP and CNN in the case of *N*_*B*_ and holograms. **(C-D)** Training curves of VGG19-PCA-MLP (C) and CNN (D). **(E-F)** Fluctuations and standard deviations of the validation accuracy (E) and the validation loss (F) during the training. * and ** indicate the statistical significance with p-value, 2.30 × 10^−107^ and 1.03 × 10^−255^ respectively by two sample F-tests. **(G)** Comparison of computing times of the feature extraction (VGG19), MLP training, and CNN training (64 epochs).

The DTL also used significantly less computational resources than the CNN. For one-time training, the VGG19-PCA-MLP took 30% less time than the CNN (**Fig. 7G**; NVIDIA GTX 1080Ti was used). In the VGG19-PCA-MLP training, the majority of time was spent in the feature extraction (VGG-PCA) rather than MLP training (**Fig. 7G**). Once the features of the training set were extracted, the repeated training was highly efficient whereas the CNN training required the feature extraction in every step of the training. Optimizing VGG19-PCA-MLP was much more efficient compared to the CNN, which could allow for training VGG19-PCA-MLP in a computation limited POC devices. Moreover, combining the automatic cell candidate identification and our DTL based prediction, it took 7.7 seconds to process the whole FOV image (3000 × 3500 pixels, the number of cell candidate: 100). In summary, these results demonstrate the feasibility of hologram classification without reconstruction, simplifying the workflow and decreasing the computational cost for a POC application.

## DISCUSSION

We have demonstrated that DTL approaches can effectively classify holograms of bead-bound cells without reconstructing original object images. The conventional reconstruction involves heavy computation, executing iterative phase recovery processes. Our DTL approach requires much less computational power, which could allow for POC devices to train and predict raw holograms.

Intriguingly, our neural networks reliably handled overlapping interference patterns among cells or between cells and unbound beads. In our training set, the target cells were positioned at the centers of the images and other cells or unbound beads were away from the image centers. More than 70% of intensity is concentrated in the first inner circle of a hologram, whereas interferences between two holograms usually happen in the fringes and have much weaker signal strength. Conceivably, the trained networks placed more weight on the hologram center, effectively ignoring fringe patterns.

Our DTL approach could offer appealing new directions to further advance LDIH: i) deep learning-based training/classification can be executed at the local device level without complex computation; ii) not relying on high-resolution reconstructed images, the classification network is robust to experimental noises such as reconstruction errors or artifacts; and iii) the network is elastic and can be continuously updated for higher accuracy in POC devices. With these merits, we envision that the developed ML networks will significantly empower LDIH, realizing a truly POC diagnostic platform.

## METHODS

### Data Collection

Samples were prepared by labeling cancer cells (SkBr3, A431) with polystyrene beads (diameter, 6 μm). We prepared four different sets of beads. Three sets were conjugated with antibodies against different molecular targets: EGFR, EpCAM, and HER2; the fourth set was conjugated with control IgG antibodies. Aliquots of cells were labeled with each set of the bead. Labeled cells, suspended in buffer, were loaded on a microscope slide, and their holograms were imaged using LDIH system^3^. To prepare the dataset set for classification, we reconstructed object images from holograms using a previously developed algorithm^3^. We cropped holograms (270 × 270 pixels) around the position of the automatically detected cell candidates (see Cell Candidate Detection below). Three researchers manually annotated the holograms of the cropped cell candidates with the following labels: the numbers (0, 1, 2, 3, ≥4) of the beads attached to cells, the beads unattached to cells, multiple cells, and artifacts. Later, we collectively labeled the beads unattached to cells, multiple cells, and artifacts as ‘background.’

### Cell Candidates Detection

We implemented computational methods which automatically localized the single-cell candidates in the hologram image based on the diffracted patterns of concentric circles in holograms^32^. The algorithm uses the fact that the gradient directions of holograms on concentric circles converge to the centers of the diffraction patterns. The detailed detection procedure is the following:

1. We normalized the holograms by dividing the pixel values by background and then rescaled them into a range [0, 255].
2. We denoised the normalized hologram using Gaussian blurring with 6-pixel size (MATLAB function *imgaussfilt()*). Then we calculated the gradient direction and magnitude of the denoised holograms using the MATLAB build-in function *imgradient ()* with ‘*prewitt*’ method.
3. We thresholded the gradient magnitude images using the threshold value 8.0, which removed the small gradient magnitude pixels and generated the binary mask. Then, the gradient direction images were masked by the gradient magnitude binary mask.
4. Along each direction in the masked gradient direction images, the frequencies of the gradient directions were accumulated within a specified range (50-pixel length, which generated the frequency maps of the gradient direction.
5. We denoised the frequency accumulation map using Gaussian blurring with 3-pixel size (MATLAB function *imgaussfilt()*). Then, we thresholded the denoised frequency accumulation map using the top 1% of the pixel values, and locate the center candidates. Then, we cropped 270 × 270 hologram and object image patches around the detected candidate center positions.

### Labelling Training Set

Three annotators independently labeled cropped holograms and their corresponding object images. In order to balance the class distribution, we augmented the image data labeled with *N*_*B*_= 1, 2, 3, and ≥4. The augmentation was performed by two strategies: rotation with a range of [0, 40] and zooming-in with the maximum value, 0.2 using Keras library.

### Machine Learning Classification

Using VGG19 pretrained model, we extracted image features from cropped holograms and object images. Since VGG19 was originally used for color images (RGB channels), and our data in a gray scale were duplicated to each channel. For data preprocessing, we perform the standard normalization, where each image patch was subtracted by its mean value and divided by its standard deviation. After the features extracted from VGG19, PCA (Principal Component Analysis) was performed to reduce the dimensionality of the data from 32768 to 500.

After the feature extraction step, we used an MLP (Multilayer Perceptron) consisting of three fully-connected neural network blocks for the classification. The first two blocks have a fully-connected (FC) layer, Batch Normalization layer, ReLU activation and Dropout layer (parameter: 0.5). The FC layers in the first two blocks have the sizes of 128 and 64, and the L2 norm regularizer (parameter: 0.05). The third block has an FC layer with ‘softmax’ activation. Also, Support Vector Machine (SVM) and Random Forest (RF) were applied to compare the performance with the MLP. The parameters of SVM and RF were optimized by the grid-search method in *sklearn* package shown below.

|  | n_estimators | max_features | max_depth |
| --- | --- | --- | --- |
| RF in binary | 20, 50, 100 | ‘Auto’, ‘sqrt’ | [none, 2, 20, 50] |
| classification |  |  |  |
| RF in 5 categories classification | 20, 50, 100, 500 | ‘Auto’, ‘sqrt’ | [none, 2, 20, 50, 100] |

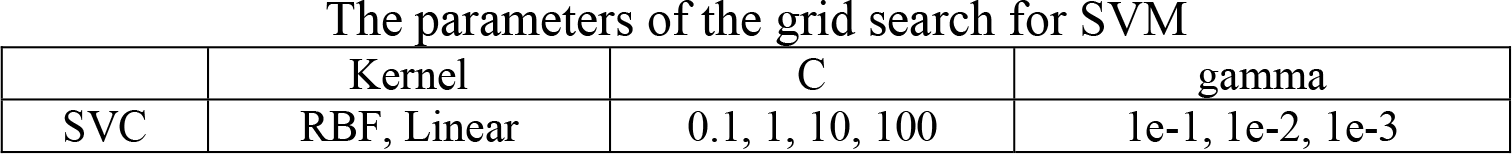

The parameters of the grid search for SVM
|  |  |  |  |
| --- | --- | --- | --- |
|  | Kernel | C | gamma |
| SVC | RBF, Linear | 0.1, 1, 10, 100 | 1e-1, 1e-2, 1e-3 |

To show the roles of the pretrained VGG19, we trained a CNN convolution neural using the same dataset (**Fig. 7A**). The CNN has three feature extraction blocks consisting of two convolutions layers and one max-pool layer (4 × 4.) After this feature extraction, the same MLP structure was used for the classification.

### Performance evaluation of the classifiers

We split the augmented dataset into three groups in a stratified fashion using the class labels: training, validation and test sets (64:16:20). The training set (64%) was used for training the network. The validation set (16%) was used for the model selections. After the training, the classification performance was evaluated using the testing set (20%). For the robust statistical analysis, we repeated the training 20 times. The performance measures were the accuracy, Cohen’s Kappa coefficient (Kappa)^34^, relative classifier information (RCI)^36^. Kappa is a standard metric for a multi-categorical classification and RCI, as an entropy-based measure, is also suitable to evaluate the performance by measure the reduced uncertainty by the classifier in comparison to the prior class distribution. For the classification involved with negative and positive bead attachment, we also measure sensitivity and specificity using *sklearn.metrics* python package. The samples of the positive bead attachment were treated as ‘positive,’ and the other cases were treated as ‘negative’. For the statistical testing, we used unpaired two-tailed Wilcoxon rank sum test method which does not rely on the assumption of the Gaussian distribution.

### Molecular Profiling

To quantify the distribution of the proportion or the frequency of the number of the attached beads (*N*_*B*_), we chose 18 images, whose cell candidates were larger than 15. For each image, we calculated the proportion and the frequency of the predicted and the actual numbers of the attached beads in each hologram.

#### Comparison between VGG19-PCA-MLP and CNN

To evaluate the performance between VGG19-PCA-MLP and CNN, the performance measures including accuracy, Kappa and RCI were used as described above. Then, the fluctuations of the validation accuracy and loss were measured as follows: we selected the last 20 epochs for each training process, and then calculate the residuals by subtracting the sample mean value. We repeated the training twenty times with random data splitting. The statistical test for the difference of variance was performed by two sample F-test.

#### Code availability statement

The code used in the current study is available from the corresponding author upon reasonable request.

#### Data availability statement

The datasets used in the current study are available from the corresponding author on reasonable request.

## Acknowledgments

We thank NVIDIA for providing us with a GPU cards (NVIDIA Hardware Grant Program, K.L.) and Microsoft for Azure cloud computing resources (Microsoft Azure Research Award, K.L.). This work is supported by the WPI Start-up Fund for new faculty. The authors were supported in part by the WPI Start-up Fund for new faculty (K.L); a generous gift by Boston Scientific (K.L); NIH grants R01-CA229777 (H.L.), U01-CA233360 (H.L.), R01-CA204019 (R.W.), R01-EB010011 (R.W.), R01-EB00462605A1 (R.W.), K99-CA201248-02 (H. I.); Liz Tilberis Award - Ovarian Cancer Research Fund (C.M.C); the Lustgarten Foundation (R.W.); and MGH Scholar Fund (H.L.).

## Author Contributions

S.K. initiated the project, designed and trained the classifiers, coordinated the collaboration as a research scientist at WPI and non-employee research personnel at MGH, and wrote the manuscript; C.W. implemented the automatic detection of cell candidates, and designed and trained the classifiers; B.Z performed training of CNNs. H.I. and J.M set up the imaging system and generated the hologram data; H.J.C, J.T. and N.C. prepared the training set and tested the ML models; C.M.C and R.W coordinated the experiments with cancer cells; K.L. and H.L. coordinated the study and wrote the final version of the manuscript and supplement. All authors discussed the results of the study.

## Author Information

Correspondence and requests for materials, code and data should be addressed to K.L. and H. L.

## REFERENCES

1 Garcia-Sucerquia, J. et al. Digital in-line holographic microscopy. Appl Opt 45, 836–850 (2006).

2 Greenbaum, A. et al. Imaging without lenses: achievements and remaining challenges of wide-field on-chip microscopy. Nat Methods 9, 889–895, doi:10.1038/nmeth.2114 (2012).

3 Im, H. et al. Digital diffraction analysis enables low-cost molecular diagnostics on a smartphone. Proc Natl Acad Sci U S A 112, 5613–5618, doi:10.1073/pnas.1501815112 (2015).

4 Xu, W., Jericho, M. H., Meinertzhagen, I. A. & Kreuzer, H. J. Digital in-line holography for biological applications. Proc Natl Acad Sci U S A 98, 11301–11305, doi:10.1073/pnas.191361398 (2001).

5 Gurkan, U. A. et al. Miniaturized lensless imaging systems for cell and microorganism visualization in point-of-care testing. Biotechnol J 6, 138–149, doi:10.1002/biot.201000427 (2011).

6 Greenbaum, A. et al. Increased space-bandwidth product in pixel super-resolved lensfree on-chip microscopy. Sci. Rep. 3, 1717, doi:10.1038/srep01717 (2013).

7 Zhu, H., Isikman, S. O., Mudanyali, O., Greenbaum, A. & Ozcan, A. Optical imaging techniques for point-of-care diagnostics. Lab Chip 13, 51–67, doi:10.1039/c2lc40864c (2013).

8 Fienup, J. Phase retrieval algorithms: a comparison. Appl Opt 21, 2758–2769, doi:10.1364/AO.21.002758 (1982).

9 Mudanyali, O., Oztoprak, C., Tseng, D., Erlinger, A. & Ozcan, A. Detection of waterborne parasites using field-portable and cost-effective lensfree microscopy. Lab Chip 10, 2419–2423, doi:10.1039/c004829a (2010).

10 Mudanyali, O. et al. Compact, light-weight and cost-effective microscope based on lensless incoherent holography for telemedicine applications. Lab Chip 10, 1417–1428, doi:10.1039/c000453g (2010).

11 Gerchberg, R. & Saxton, W. A practical algorithm for the determination of phase from image and diffraction plane pictures. SPIE milestone series MS 93, 306–306 (1994).

12 Fienup, J. R. Reconstruction of an object from the modulus of its Fourier transform. Optics Letters 3, 27–29 (1978).

13 Latychevskaia, T. & Fink, H.-W. Solution to the twin image problem in holography. Physical Review Letters 98, 233901 (2007).

14 Rivenson, Y., Zhang, Y., Gunaydin, H., Teng, D. & Ozcan, A. Phase recovery and holographic image reconstruction using deep learning in neural networks. Light: Science & Applications, doi:10.1038/lsa.2017.141 (2017).

15 Shen, D., Wu, G. & Suk, H.-I. Deep learning in medical image analysis. Annual Review of Biomedical Engineering 19, 221–248 (2017).

16 Esteva, A. et al. Dermatologist-level classification of skin cancer with deep neural networks. Nature 542, 115–118, doi:10.1038/nature21056 (2017).

17 Gulshan, V. et al. Development and Validation of a Deep Learning Algorithm for Detection of Diabetic Retinopathy in Retinal Fundus Photographs. JAMA 316, 2402–2410, doi:10.1001/jama.2016.17216 (2016).

18 Ehteshami Bejnordi, B. et al. Diagnostic Assessment of Deep Learning Algorithms for Detection of Lymph Node Metastases in Women With Breast Cancer. JAMA 318, 2199–2210, doi:10.1001/jama.2017.14585 (2017).

19 Pratt, L. Y. Discriminability-based transfer between neural networks. Advances in Neural Information Processing Systems. 204–211 (1993).

20 Yosinski, J., Clune, J., Bengio, Y. & Lipson, H. How transferable are features in deep neural networks? Advances in Neural Information Processing Systems. 3320–3328 (2014).

21 Razavian, A. S., Azizpour, H., Sullivan, J. & Carlsson, S. CNN features off-the-shelf: an astounding baseline for recognition, Computer Vision and Pattern Recognition Workshops (CVPRW), 2014 IEEE Conference on. 512–519 (IEEE) (2014).

22 Donahue, J. et al. Decaf: A deep convolutional activation feature for generic visual recognition, International Conference on Machine Learning. 647–655 (2014).

23 Oquab, M., Bottou, L., Laptev, I. & Sivic, J. Learning and transferring mid-level image representations using convolutional neural networks. Computer Vision and Pattern Recognition (CVPR), 2014 IEEE Conference on. 1717–1724 (IEEE) (2014).

24 Zeiler, M. D. & Fergus, R. Visualizing and understanding convolutional networks. European Conference on Computer Vision. 818–833 (Springer) (2014).

25 Choi, J. Y. et al. Multi-categorical deep learning neural network to classify retinal images: A pilot study employing small database. PLoS One 12, e0187336, doi:10.1371/journal.pone.0187336 (2017).

26 LeCun, Y. et al. Handwritten digit recognition with a back-propagation network, Advances in Neural Information Processing Systems, 396–404 (1990).

27 LeCun, Y., Bengio, Y. & Hinton, G. Deep learning. Nature 521, 436–444, doi:10.1038/nature14539 (2015).

28 Krizhevsky, A., Sutskever, I. & Hinton, G. E. Imagenet classification with deep convolutional neural networks. Advances in Neural Information Processing Systems. 1097–1105 (2012).

29 Simonyan, K. & Zisserman, A. Very deep convolutional networks for large-scale image recognition. International Conference on Learning Representations (2015).

30 Deng, J. et al. Imagenet: A large-scale hierarchical image database, Computer Vision and Pattern Recognition, 2009. CVPR 2009. IEEE Conference on. 248–255 (IEEE) (2009).

31 Fung, J. et al. Measuring translational, rotational, and vibrational dynamics in colloids with digital holographic microscopy. Opt Express 19, 8051–8065, doi:10.1364/OE.19.008051 (2011).

32 Cheong, F. C. et al. Flow visualization and flow cytometry with holographic video microscopy. Optics Express 17, 13071–13079 (2009).

33 Maaten, L. v. d. & Hinton, G. Visualizing data using t-SNE. Journal of Machine Learning Research 9, 2579–2605 (2008).

34 Cohen, J. A coefficient of agreement for nominal scales. Educational and Psychological Measurement 20, 37–46 (1960).

35 Statnikov, A., Wang, L. & Aliferis, C. F. A comprehensive comparison of random forests and support vector machines for microarray-based cancer classification. BMC Bioinformatics 9, 319, doi:10.1186/1471-2105-9-319 (2008).

36 Bhattacharyya, P., Sindhwani, V. & Rakshit, S. Information Theoretic Feature Crediting in Multiclass Support Vector Machines, Proceedings of the First SIAM International Conference on Data Mining. 1–18 (2001).

